# Towards a unified theory of the reference frame of the ventriloquism aftereffect

**DOI:** 10.1101/2021.03.31.437664

**Authors:** Peter Lokša, Norbert Kopčo

## Abstract

The ventriloquism aftereffect (VAE), observed as a shift in the perceived locations of sounds after audio-visual stimulation, requires reference frame alignment since hearing and vision encode space in different reference frames (head-centered vs. eye-centered). Previous experimental studies reported inconsistent results, observing either a mixture of head-centered and eye-centered frames, or a predominantly head-centered frame. Here, a computational model is introduced to examine these inconsistencies. Based on experimental data, the model uses the measured size of the ventriloquism effect to predict the VAE adaptation in the auditory spatial map. Versions of the model examine whether the adaptation is induced by visual signals in head-centered frame, eye-centered frame, by eye-gaze direction-dependent signals, or their combination, and whether some biases are induced by the saccade-to-auditory-target response method used in the experiments. The model is first evaluated on three separate data sets. It can predict them well even without explicit need for an eye-centered signals influencing VAE, suggesting that the reference frame of VAE is mainly head-centered. The model predictions are qualitatively similar but less accurate when all three data sets are combined, suggesting that interactions between individual neural mechanisms are more complex than the simple linear combination assumed in the model.

## 1. Introduction

Auditory spatial perception is highly adaptive and visual signals often guide this adaptation. In the “ventriloquism aftereffect” (VAE), the perceived location of sounds presented alone is shifted after repeated presentations of spatially mismatched visual and auditory stimuli (Bertelson et al., 2006, Recanzone, 1998, Woods and Recanzone, 2004). Complex transformations of spatial representations in the brain are necessary for the visual calibration of auditory space to function correctly, as visual and auditory spatial representations differ in many important ways (van Opstal, 2016). Here, we propose a computational model to examine the visually guided adaptation of auditory spatial representation in the VAE, the related transformations between the reference frames (RFs) of auditory and visual spatial encoding, and the effect that the saccade response method, used in many RF of VAE studies, might have on the results.

We primarily examine the RF in which the VAE occurs. While visual space is initially encoded relative to the direction of eye gaze, the cues for auditory space are computed relative to the head orientation (Groh and Sparks, 1992). A means of aligning these RFs is necessary by the stage at which the visual signals guide auditory spatial adaptation. Several models were developed to describe the ventriloquism aftereffect in humans and birds. The bird models predict the VAE in the barn owls (Haessly et al., 1995, Oess et al., 2020) which cannot move their eyes and therefore their auditory and visual RFs are aligned. The human models mainly focused on spatial and temporal aspects of the ventriloquism aftereffect (Beierholm et al., 2009, Bosen et al., 2018, Kording et al., 2007, Odegaard et al., 2015, Shinn-Cunningham et al., 2005, Watson et al., 2019, Wozny et al., 2010), not considering the different RFs. There are models of the audio-visual reference frame alignment, but those only consider audio-visual integration (Odegaard et al., 2019, Razavi et al., 2007) and multi-sensory integration (Pouget et al., 2002) when the auditory and visual stimuli are presented simultaneously, not the adaptation and transformations underlying the VAE.

Our experimental studies examining RF of VAE in humans and monkeys provided inconsistent results (described in detail in Section 2). A mixture of eye-centered and head-centered RFs was identified for recalibration locally induced in the central region of the audio-visual field (Kopco et al., 2009) while the head-centered RF dominated when was VAE induced in the audio-visual periphery (Kopco et al., 2019). Additionally, a recent study in which the VAE was induced over a wide spatial area also concluded that the RFs are mixed (Watson, 2021). These results imply that the RF used in the VAE is dependent on the region in which the VAE is induced, possibly due to a non-homogeneity in the auditory spatial representation. Specifically, recent evidence suggests that, in mammals, auditory space encoding is based on two or more spatial channels roughly aligned with the left and right hemifields of the horizontal plane (Groh, 2014, Grothe et al., 2010), implying that the spatial representation’s adaptability might be spatially non-uniform. For example, it might be more locally adaptable in the subregions of space covered by both hemispheric channels, i.e., in the center, than in the regions dominated by one channel (periphery). Our current modeling focuses on a simpler alternative in which the RF transformations are the same across the audio-visual field, and the observed region-dependence of the RF is due to saccades to auditory targets, as saccades, which are eye-centered, were used to measure VAE in our previous studies.

A secondary goal of the current model to propose a mechanism to describe a new adaptive phenomenon observed in the ventriloquism study of Kopco et al. (2019) (again, described in detail in Section 2). In that study, adaptation was unexpectedly induced by spatially aligned audio-visual stimuli, while no such adaptation was observed in Kopco et al. (2009).

Here, we first summarize the experimental data from Kopco et al. (2009, 2019) to explain the modeled phenomena (Section 2). Then, the model is introduced (Section 3) and evaluated (Sections 4 and 5).

## 2. Summary of data of Kopco et al. (2009, 2019)

In two separate experiments, ventriloquism was induced by audio-visual training trials either in the central (Kopco et al., 2009) or peripheral (Kopco et al., 2019) subregion of the horizontal audio-visual field while the eyes fixated one fixation point (FP; red ‘+’ symbol; upper and middle panels of Figure 1A, respectively). The audio-visual stimuli consisted of a sound paired with an LED. The aftereffect was evaluated on interleaved auditory-only (A-only) probe trials using a wide range of target locations while the eyes fixated one of two FPs (lower panel of Figure 1A). The listener’s task in both audio-visual and auditory-only trials was to perform a saccade to the perceived location of the auditory stimulus from the FP.

**Figure 1:**
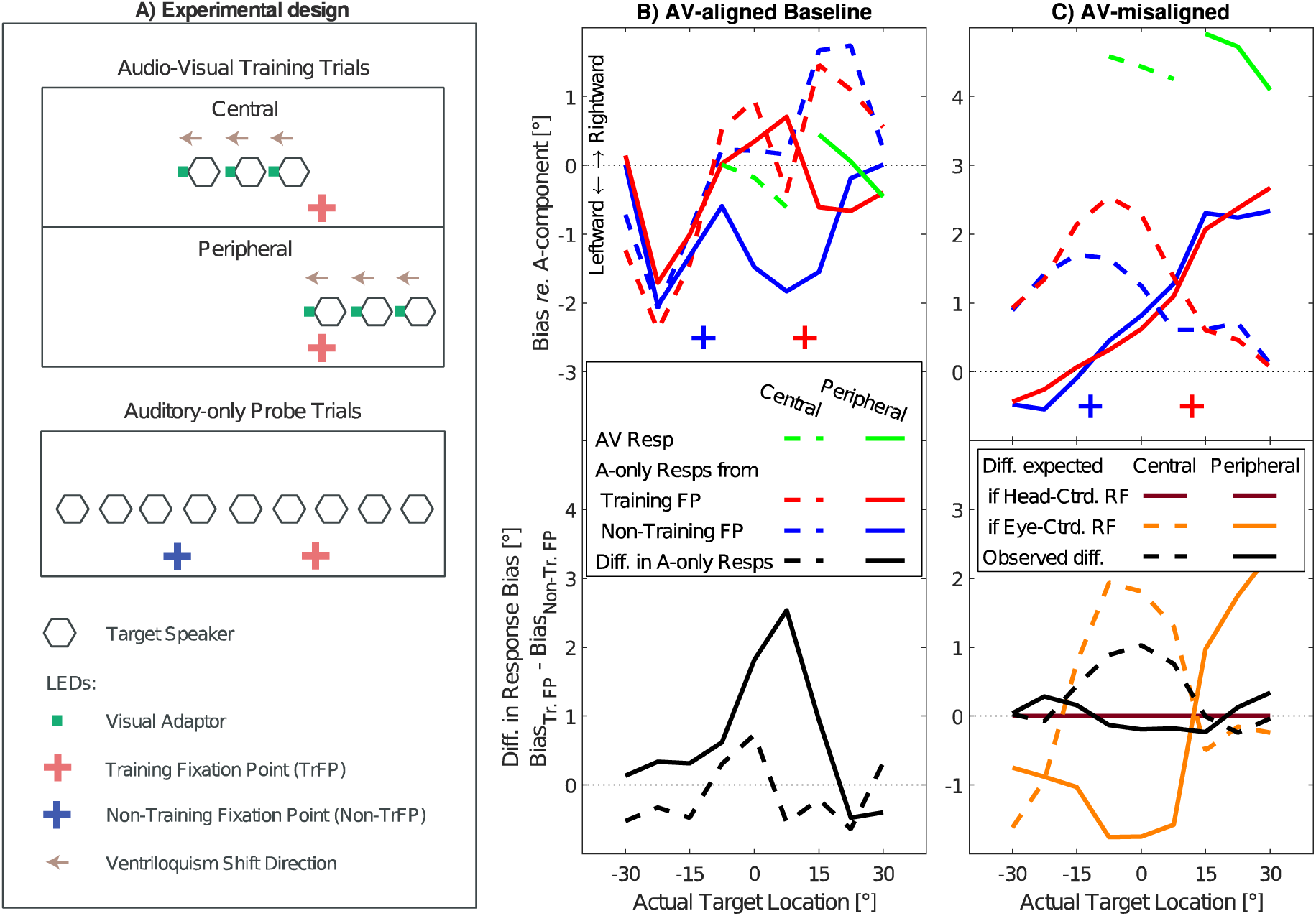
Experimental design and results from Kopco et al. (2009; 2019). A) Experimental design: nine loudspeakers were evenly distributed at azimuths from −30° to 30°. Two fixation points (FP) were located 10° below the loudspeakers at ±11.75° from the center. On training trials, audio-visual stimuli were presented either from the central region (N. Kopco et al., 2009) or peripheral region (N. Kopco et al., 2019), while the subject fixated the red FP. The audio-visual (AV) stimuli consisted of a sound paired with an LED offset by −5°, 0°, or −5° (offset direction fixed within a session). On probe trials, the sound was presented from any of the loudspeakers while the eyes fixated one of the FPs. B) AV-aligned results: Across-subject mean biases in AV (green) and A-only trials (red and blue lines for respective FPs), corresponding, respectively, to the ventriloquism effect and aftereffect from the runs with AV-aligned stimuli. Black lines show the differences between the respective red and blue lines, i.e., differences between the biases found for the two FPs. C) AV-misaligned results: green, red, blue and black lines as in the AV-aligned results. Brown and beige lines show the expected results for the difference data if the RF is, respectively, head-centered and eye-centered. Note: Error bars have been omitted for clarity. N=7 in both experiments. All horizontal axes are plotted in head-centered reference frame.

The responses to AV stimuli were always very near the visual components of the AV stimuli, both in the AV-aligned baseline (green dashed and solid lines in Figure 1B corresponding, respectively, to the peripheral and central experiments) and in the conditions with the visual component shifted (green lines in Figure 1C). The displaced V component in the AV-misaligned conditions induced a local ventriloquism aftereffect when measured with the eyes fixating the training FP (the red solid and dashed lines in Figure 1C show that maximum ventriloquism was always induced in the trained subregion of the auditory space). The critical manipulation of these experiments was that a half of probe trials was performed with eyes fixating a new, non-training FP (blue ‘+’ symbol), located 23.5° to the left of the training FP (red ‘+’ symbol at +11.75°). If the RF of the VAE is purely head-centered, then moving the eyes to the new FP location for the testing was expected to have no effect, resulting in the same pattern of ventriloquism for the non-training and training FPs. On the other hand, if the RF is purely eye-centered, the observed pattern of induced shifts was expected to move with the eyes by the 23.5° when the probe trial is performed from the new, non-trained fixation. The experimental data showed that, in the central experiment, moving the fixation resulted in a smaller ventriloquism aftereffect with the peak moving in the direction of eye gaze (blue vs. red dashed line), while in the peripheral experiment only a negligible effect of the eye fixation position was observed (blue vs. red solid line).

To better visualize these results, the lower panel of Figure 1C shows data expressed as difference between responses from training vs. non-training FPs from the respective upper panels, along with the expected patterns of results for the two RFs. The head-centered RF always predicts that the effect would be identical for the two FPs. Thus, all head-centered differences (brown lines) are expected at zero. The solid and dashed yellow lines show, respectively, for the peripheral and central data, the eye-centered RF expected patterns obtained by subtracting from the red lines the same red lines shifted 23.5° to the left. Finally, the black solid and dotted lines show the actual differences between the respective red and blue data from the upper panel. For the central data, the black dashed line falls approximately in the middle between the head-centered and eye-centered predictions, showing a mixed nature of the RF of the VAE induced in this region. On the other hand, the black solid line is always near zero, showing that the RF of the VAE induced in the periphery is predominantly head-centered. The main goal of the current modeling is to examine this inconsistency.

While the results in Figure 1C are based on the ventriloquism aftereffect induced by AV-misaligned stimuli, Figure 1B shows the baseline data obtained in runs with auditory and visual stimuli aligned. In the central-training experiment, the responses from the two FPs were similar, unbiased at the central locations and with a slight expansive bias in the periphery (both red and blue dotted lines are near zero in the center, negative in the left-hand portion and positive in the right-hand portion of the graph). On the other hand, in the peripheral-training experiment the responses for the middle three targets differed between the two fixations, where the non-training FP responses fell well below the training-FP responses (compare the red and blue solid lines). Thus, the peripheral AV-aligned stimuli induced a fixation-dependent adaptation in the auditory-only responses in the central region, an adaptation phenomenon that has not been previously described. The black dashed and solid lines in Figure 1B showing the difference between the corresponding data from the upper panel, highlight the FP-dependence of the peripheral data in contrast to the FP-independence in the central data. The secondary goal of the current modeling is to propose a mechanism to explain this result.

## 3. Model Description

### 3.1 Overview

Figure 2A shows the outline of the model. The model predicts the ventriloquism aftereffect (i.e., the azimuthal bias in a response to an auditory-only probe; red and blue lines in Figure 1B-C) as a function of the A-only probe azimuth and the FP location (the “Response” vs. “Probe stimulus and FP” blocks in Figure 2A). To generate the prediction, it only uses information about the training AV stimulus locations and the measured AV response biases for those locations (i.e., the ventriloquism effect; green lines in Figure 1B-C) for a given type of AV stimuli (“Ventriloquism” block in Figure 2A). The prediction is computed as an additive combination of two components (the “sum” operation). The main component is the “Auditory space representation” block (Bosen et al., 2018) which is locally adapted by the audio-visual ventriloquism signals in different reference frames (“HC” and “EC” arrows) and which can also be influenced by information about eye position (“FP-dependent attenuation”). The second component, the “Saccade-related bias” block, represents the saccade response bias in eye-centered reference frame, which is not directly adapted by ventriloquism and which is assumed to be present *a priori* when saccades are used as a response method. Finally, it is assumed that there are interactions between the saccade-related and ventriloquism effects (the “Saccade-related bias” block is linked to the “Auditory space representation” block).

**Figure 2:**
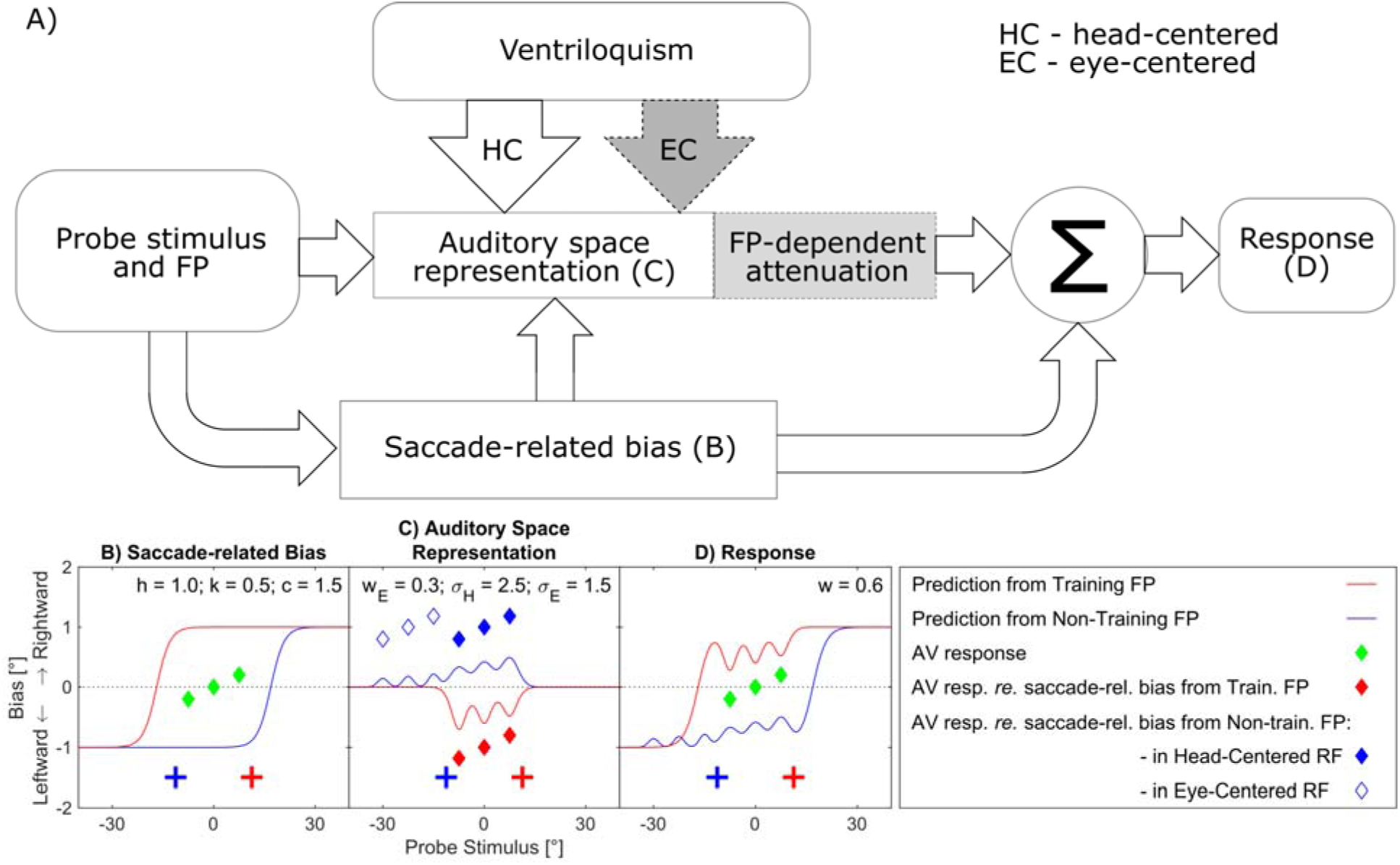
Structure of the model in its 4 versions and illustration of its operation. A) Block diagram of the model in which optional model components that differentiate the 4 model versions are shown in gray. The model predicts the response bias (“Response”) as a function of the “Probe stimulus location and FP”, with additional input parameters such as the training locations and the observed ventriloquism effect at the training locations (“Ventriloquism”). The prediction is determined as a sum of two components (square blocks). Component 1: “Auditory space representation” is adapted by AV signals either only in HC reference frame (used in the HC model version; “HC” arrow) or in a combination of HC and EC RFs (used in the HEC model; “HC” arrow and the optional “EC” arrow). In the dHC and dHEC model versions, the size of the auditory space adaptation is reduced depending on the distance of the current FP from the training FP (“FP-dependent attenuation” optional subblock). Component 2: “Saccade-related bias” observed when saccades are used for responses. Labels (B), (C), and (D) within blocks refer to respective panels below that illustrate the function of the blocks by showing the outputs of the model components. Panels B-D: An illustrative simulation of the HEC model version for AV stimulus locations and responses shown by green diamonds in panel B (i.e., both near 0). B) Saccade-related bias predictions (red and blue lines) for the two fixation points (crosses). C) Adaptation of the auditory space representation resulting from the saccade-related bias and AV response bias from panel B. Diamonds represent the disparity between AV response bias and saccade-related bias for the training FP (red), and non-training FP in HC RF (blue filled diamonds) and in EC RF (blue open diamonds). Lines represent predictions of auditory space adaptation induced by these disparities. (In the HC model version, there would be no open blue diamonds, and in the dHC and dHEC model versions, the blue line would be scaled down by the value of *d_f_*.) D) Response bias predicted by the model as a weighted combination of biases from panels B and C. Model parameters used for the predictions of respective model components are shown in each panel. All horizontal axes are plotted in HC RF.

The current model assumes that the auditory space representation uses a uniform population code (Carlile et al., 2001), not a hemispecific space encoding (Lingner, 2018). It describes the VAE adaptation by only considering the locations from which the auditory components of audio-visual stimuli were presented during training. Then, it assumes that the induced VAE at those locations is directly proportional to the measured ventriloquism effect and those locations, and that it decreases with distance from those locations. Thus, the model does not require input information about the direction of audio-visual stimulus displacement during training (whether the visual stimuli were shifted to the left, right, or aligned with the auditory stimuli). Instead, it only uses the information about where the training occurred and what the resulting ventriloquism effect was, and it assumes that there is a direct relation between the observed ventriloquism effect and aftereffect. Supporting this assumption, a comparison of the VE and VAE data at the trained locations (corresponding green and red lines in Figure 1C) shows that the VAE is approximately a half of VE in the experimental data.

There are four versions of the model, differing by whether they include optional components shown in gray in Figure 2A (“EC” arrow and “FP-dependent attenuation” subblock), which represent two different hypotheses about how EC signals might influence the RF of VAE. The basic version of the model, referred to as “HC model”, does not include either of the optional components. Thus, it predicts that the ventriloquism signals influencing the spatial auditory representation are purely head-centered (the “HC” arrow in Figure 2A) and all the eye-centered contributions to the observed RF of VAE come from the saccade-related biases.

In the “HEC model” version, the visual signals adapt the auditory spatial representation in both head-centered and eye-centered RFs (the optional “EC” arrow) such that the relative contribution of the HC and EC RFs can be arbitrary. Therefore, the HEC model reduces to the HC model if the weight of the EC path is set to zero, or it can produce predictions using only EC RF if the HC path weight is set to zero. Note that a purely EC-based version of the model was not considered as 1) the peripheral-experiment data only are consistent with a HC reference frame, so the model would clearly fail, and 2) the HEC model could behave as such EC-only model by appropriately adjusting the relative weight, if that were the best fit to the data.

The “dHC model” version assumes that the adaptation of spatial representation induced by ventriloquism is attenuated when the eyes shift to a new FP away from the training FP (“FP-dependent” attenuation sub-component), and that this attenuation is proportional to the distance between the training and new FP. This mechanism might be related to FP-dependent biases observed in sound localization (Lewald & Ehrenstein, 1998; Razavi et al., 2007), and it does not require saccades to be used for responses. Since this attenuation is dependent on the fixation location, it is also in the eye-centered reference frame.

Finally, the “dHEC model” incorporates both optional components and thus it assumes that both EC-referenced mechanisms described in the HEC and dHC models affect performance.

### 3.2 Detailed Specification

The following model specification applies to the most general dHEC model version, with the differences applying to the reduced versions described as needed (all variables in the model use the head-center representation and are in the units of degrees, unless specified otherwise). Panels B-D of Figure 2 provide visualizations of the behavior of the individual model components.

Equation 1 describes the predicted bias in responses *r̂* to a given auditory stimulus at location and for eyes fixating the location *f*, as a weighted sum of a saccade-related bias *r_E_* and a ventriloquism-related adaptation in auditory spatial representation *r_V_*

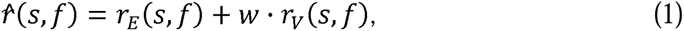

Where *w* ∈ [0, ∞] is a free parameter specifying the relative weight of the ventriloquism adaptation. The prediction is determined by considering the observed biases in AV stimulus responses *r_AV_* at the training AV stimulus locations *S_AV_* (as described below in Eq. 3).

The saccade-related bias at a specific location *x* for eyes fixating the location *f* is modeled as a sigmoidal function

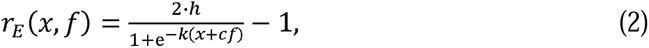

where *h*, *k*, and *c* are free parameters characterizing the sigmoid. The saccade-related bias (Figure 2B) is broad and referenced to the FP (i.e., it uses EC RF), exhibiting a combination of underestimations and overestimations commonly observed in studies of saccades to auditory targets (Gabriel et al., 2010, Razavi et al., 2007, Yao and Peck, 1997). While similar functions were previously used to model visually induced spatial auditory adaptation (Zwiers et al., 2003), the specific shape of the functions used here was chosen to be consistent with results of Gabriel et al. (2010), which observed both underestimation and overestimation in saccade responses depending on the target location (or, it can be a result of underestimation of saccade responses (Yao and Peck, 1997) combined with overestimation in peripheral auditory localization estimates with a centrally fixed eyes (Bosen et al., 2018)). The resulting predictions roughly follow the values observed in the peripheral and central no-shift data in the current study (Figure 1B).

The ventriloquism-driven auditory space adaptation causes bias defined at location *x*, for eyes fixating the location *f*, and for ventriloquism induced at training locations *S_AV_* Scausing AV response biases *r_AV_*, as:

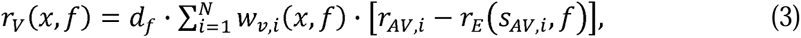

where *d_f_* is the FP-dependent attenuation of the aftereffect in the dHC and dHEC models (Eq. 8), is the number of training locations (*N* = 3 for the current study), *i* is an index through these locations, *S_AV,i_* is the *i*-th training location azimuth, and *r_AV,i_* is the AV response bias observed at the *i*-th training location (the green diamonds in Figure 2B represent the values of *r_AV,i_* in an example simulation in which these values are around 0 and in which *S_AV,i_* are at −7.5, 0, and +7.5°). The differences *r_AV,i_* − *r_E_* (*S_AV,i_*) represent the disparity between the AV response biases (green diamonds) and the saccade-related bias (red/blue lines in Figure 2B) at the training locations (where the *r_E_* value explicitly incorporates an interaction between the ventriloquism and the saccade components, in addition to the ventriloquism effect). The differences are shown in Figure 2C by the red and blue filled diamonds. *W_v,i_*(*x*) is the strength with which the disparity at the *i*-th training location adapts the spatial representation at the location *x*. In the HEC and dHEC models, this value is a weighted sum of the adaptation strengths in head-centered and eye-centered reference frames, defined as:

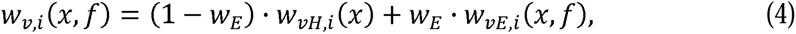

where *w_E_* ∈ (0,1) is a parameter determining the relative weight of the EC reference frame vs. the HC RF (in the HC and dHC models, *w_E_* = 0). Weights *w_vH,i_* and *w_vE,i_* use normalized Gaussian functions centered at the training locations as a measure of the influence of *i*-th training location on target at location *x*:

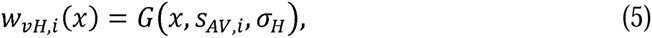

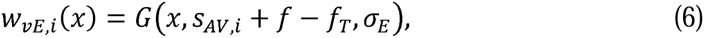

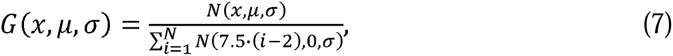

In Eqs. 5 and 6, the parameters σ*_H_* and σ*_E_* represent the width of the influence of the ventriloquism shift at individual training locations, respectively, for the two reference frames. *w_vH,i_* (Eq. 5) is always centered on the *i*-th training location in the HC RF, whereas *w_vE,i_* (Eq. 6) is centered on the *i*-th training location in the EC RF. *f_T_* is the training fixation location, equal to 11.75° in the current study (when *f* = *f_T_*, the two RFs are aligned). Finally, the Gaussian functions are normalized (Eq. 7) such that the maximum *w_vH,i_* or *w_vE,i_* after summing across the three training locations is 1 (the normalization locations 7.5 · (*i* − 2) are specific for the current training as 7.5° is the separation between the loudspeakers used in the experiments; they need to be modified to model data from experiments using other training locations).

The FP-dependent attenuation of the aftereffect from the non-training FP is defined as

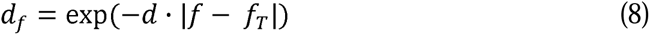

where the free parameter *d* is the rate of attenuation with distance from the current fixation *f* to the training fixation *f_T_* (in the HC and HEC models, *d* = 0). Note that the exact form of this dependence cannot be evaluated for the current data as only one training and one non-training fixation were evaluated (i.e., |*f* − *f_T_*| only had values of 0 or 23.5°).

Figure 2C shows the operation of the ventriloquism adaptation. As mentioned above, the red and blue filled diamonds are the disparities at the individual training locations driving the adaptation in HC RF. The blue open diamonds are identical to the blue filled diamonds except that they are shifted to the left by the difference between the two FPs to illustrate how the eye gaze shift affects where the adaptation is expected to occur in the EC RF. The red and blue lines are then the resulting biases *r_V_* for the two fixation locations, each corresponding to the sum of Gaussians centered at different training locations in the two RFs (and with widths defined by the σ’s). Parameter *w_E_* determines the relative weights of the peaks in the portion of blue line corresponding to the open diamonds vs. the portion corresponding to the filled diamonds. Finally, parameter *d_f_* determines how much the blue line deviation from 0 is reduced compared to the red line (i.e., how much the aftereffect is attenuated at a new fixation location). So, the blue and red lines show how visually guided adaptation is local and RF-dependent, decreasing with distance from the AV stimulus locations in HC and EC RFs. It also shows that since adaptation causes shifts from the saccade-bias response locations towards AV response locations, if AV responses fall on saccade bias locations, no visually guided adaptation is predicted to occur. Finally, Figure 2D shows that the model prediction is a sum of the saccade bias (from Figure 2B) and ventriloquism bias (Figure 2C) weighted by the parameter.

## 4. Simulation Methods

### 4.1 Stimuli

In the studies of Kopco et al. (2009, 2019), ventriloquism was induced by presenting AV training stimuli with visual component either shifted to the left of the auditory component (negative shift), to the right of it (positive shift), or aligned with it (no shift), while the eyes fixated one location (Figure 1A; upper and middle panels). Thus, nominally, there were 6 AV conditions (3 shift directions by 2 training regions). For these conditions, predictions could be compared to data for 9 A-only probe locations and 2 FPs (Figure 1A; lower panel). However, the main experimental results simulated here were observed after performing two subtractions: 1) subtracting the training-FP from the non-training-FP data, and 2) subtracting the positive-shift data from negative-shift data and halving the result, which gave the average aftereffect magnitude differences (black lines in Figure 1C). These “double differential” (“positive – negative” difference of “training FP – non-training FP” difference) data were the most stable as they eliminated a lot of between-subject variability related to individual biases in responses (as will be illustrated later). Therefore, to evaluate the model on these key differences, the data were also transformed into the difference representation in two steps, resulting in representation that is equivalent to the difference plots in the bottom panels of Figure 1B and C (Footnote 1).

The complete data set consisted of 108 data points [9 location x 2 (transformed) FPs x 3 (transformed) shifts x 2 training regions]. Across-subject mean and standard deviation data were used in the simulations. The model is fitted to the auditory-only trial responses, while the AV responses are used by the model as parameters for different conditions.

### 4.2 Model Fitting and Evaluation

In the simulations, the four models’ performance was compared on different subsets of the data. The basic rationale behind these comparisons was that if the minimal HC model predicts the data well, then there is no evidence that eye-centered signals affect the ventriloquism aftereffect, and all the observed eye-dependent effects can be ascribed to the saccade-related bias. On the other hand, if one of the HEC, dHC, and dHEC models performs best, it is evidence that, even after accounting for the saccade-related effects, the auditory spatial representation receives some eye-centered signals, causing adaptation that always depends on the position of the stimuli relative to the eye gaze direction.

Each simulation was performed by fitting the four models to the corresponding subset of the transformed data using a two-step procedure. First, a systematic search through the parameter space was performed, using all combinations of 10 values for each parameter, listed in Table 1 (HC model used 5 parameters, dHC 6, HEC 7, and dHEC 8 parameters). The ranges were chosen by piloting to cover the expected behavior of the model in different conditions. Second, the best 100 parameter combinations in terms of weighted MSE were used as starting positions for non-linear iterative least-squares fitting procedure (Matlab function lsqnonlin) which, again, minimized the weighted MSE. The parameter values for the best of these fits were chosen as the optimal values listed in Table 2 and used in the result figures.

**Table 1:** Range and increments of free parameters used in systematic search through the parameter space during model simulations. Ten values of each parameter were considered with either linear or quadratic spacing.

| Parameter | Range |  | Increments |
| --- | --- | --- | --- |
|  | min | max |  |
| $h, w$ | 0 | 2 | linear |
| $k$ | 0.01 | 20 | quadratic |
| $c$ | 0 | 1.5 | quadratic |
| $w_E$ | 0 | 1 | linear |
| $\sigma_H, \sigma_E$ | 1 | 20 | linear |
| $d_f$ | 0 | 1 | linear |

**Table 2:** Fitted model parameters and model performance for each simulation. AICc and MSE were calculated on the data used in a given simulation. ΔAIC is the increase in AICc for a given model *re* the model with the lowest AICc. The underlined model names indicate the model version with substantial evidence of better fit to the data (i.e., rounded up ΔAIC smaller than 2).

| Simulation | Model | Fitted parameter values |  |  |  |  |  |  |  | Performance |  |  |
| --- | --- | --- | --- | --- | --- | --- | --- | --- | --- | --- | --- | --- |
| | | $h$ | $k$ | $c$ | $w$ | $w_E$ | $\sigma_H$ | $\sigma_E$ | $d_f$ | AICc | ΔAIC | MSE |
| No Shift<br>(Figure 3) | <u>HC</u> | 1.03 | 0.31 | 1.14 | 1.01 | - | 12.06 | - | - | 130.9 | 2.4 | 1.59 |
|  | <u>HEC</u> | 1.13 | 0.17 | 0.95 | 1.24 | 0.36 | 12.84 | 2.98 | - | 128.5 | - | 1.26 |
|  | dHC | 1.03 | 0.31 | 1.14 | 1.01 | - | 12.06 | - | 1.00 | 133.8 | 5.3 | 1.59 |
|  | dHEC | 1.13 | 0.17 | 0.95 | 1.24 | 0.36 | 12.84 | 2.98 | 1.00 | 131.9 | 3.3 | 1.26 |
| Central<br>(Figure 4A) | HC | 1.01 | 5.64 | 0.67 | 0.40 | - | 18.79 | - | - | 176.2 | 15.6 | 5.48 |
|  | HEC | 0.96 | 5.60 | 0.67 | 0.48 | 0.30 | 18.14 | 5.01 | - | 170.2 | 9.6 | 3.86 |
|  | dHC | 1.01 | 6.84 | 0.67 | 0.52 | - | 15.20 | - | 0.68 | 160.6 | - | 3.22 |
|  | dHEC | 0.96 | 13.46 | 0.67 | 0.51 | 0.17 | 17.99 | 2.65 | 0.74 | 162.0 | 1.4 | 2.74 |
| Peripheral<br>(Figure 4B) | <u>HC</u> | 0.83 | 3.40 | 1.33 | 0.55 | - | 12.43 | - | - | 136.3 | - | 1.73 |
|  | HEC | 0.82 | 5.33 | 1.33 | 0.56 | 0.04 | 12.12 | 4.91 | - | 141.9 | 5.6 | 1.68 |
|  | dHC | 0.83 | 3.27 | 1.33 | 0.55 | - | 12.43 | - | 1.00 | 139.1 | 2.8 | 1.73 |
|  | dHEC | 0.82 | 2.85 | 1.33 | 0.56 | 0.04 | 12.12 | 4.91 | 1.00 | 144.2 | 7.9 | 1.68 |
| All Data<br>(Figure 5) | HC | 0.79 | 0.82 | 1.15 | 0.49 | - | 14.21 | - | - | 444.7 | 10.5 | 3.25 |
|  | HEC | 0.77 | 0.76 | 1.13 | 0.53 | 0.15 | 13.35 | 4.83 | - | 436.9 | 2.7 | 2.89 |
|  | dHC | 0.79 | 0.91 | 1.17 | 0.54 | - | 13.74 | - | 0.85 | 436.4 | 2.2 | 2.95 |
|  | dHEC | 0.77 | 0.82 | 1.15 | 0.55 | 0.11 | 13.64 | 4.36 | 0.90 | 434.2 | - | 2.76 |

To compare the models’ performance while accounting for the number of parameters used by each model, we computed the Akaike information criterion AICc (Burnham and Anderson, 2004, Taboga, 2017) for each optimal fit, defined as:

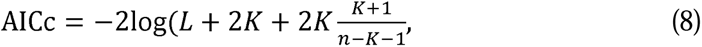

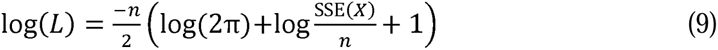

where *n* is the number of experimental data points, *K* is the number of fitted parameters, and SSE(*X*) is the sum of squares of errors across the data points (i.e., differences between predictions and across-subject mean data *x*_i_) weighted for each data point by the inverse of its across-subject standard deviation 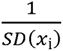. In general, the model with the lower AICc is considered to be a better fit for the data. We use the rule that the model with the lower AICc is substantially better than an alternative model only if the rounded up value ΔAIC is larger than 2.

## 5. Results

Four model evaluations were performed, each assessing the four model variants on a different subset of the Kopco et al. (2009, 2019) data. No Shift evaluation assessed the models on the AV-aligned no-shift data from both experiments (Figure 1B), while in the Central Data and Peripheral Data evaluations, the respective data from the AV-misaligned conditions in the two experiments (dashed and solid lines in Figure 1C) were fitted separately. Finally, in the Combined Data evaluation the models were fitted on the complete dataset from both experiments (Figure 1B and C). The results of the 4 evaluations are summarized in Table 2, which shows for each simulation and model the fitted parameter values and the model’s performance measured using the AICc criterion and the weighted MSE.

### 5.1 No-shift data

This evaluation focused on the AV-aligned data, examining the hypothesis that *the saccade-related bias combined with auditory space representation adapted in HC RF are sufficient to explain the training-region-dependent differences in the AV-aligned data* (Figure 1B). I.e., it was predicted that the HC model can accurately describe the baseline data, and that the optional model components do not need to be considered to explain the different adaptation effects observed in central vs. peripheral AV-aligned training.

Figure 3 presents the results of the model evaluation and illustrates the function of individual model components. Panels A and B show the data (circles) and predictions of the models (lines) with the fitted parameters as listed in Table 2. (Note that only the HC and HEC models are shown and discussed, as performance was identical for the other two models). The upper row shows the main results for the difference between the FPs (identical to the black lines from Figure 1B, now with error bars included), while the middle and bottom rows show the results separately for the two FPs.

**Figure 3:**
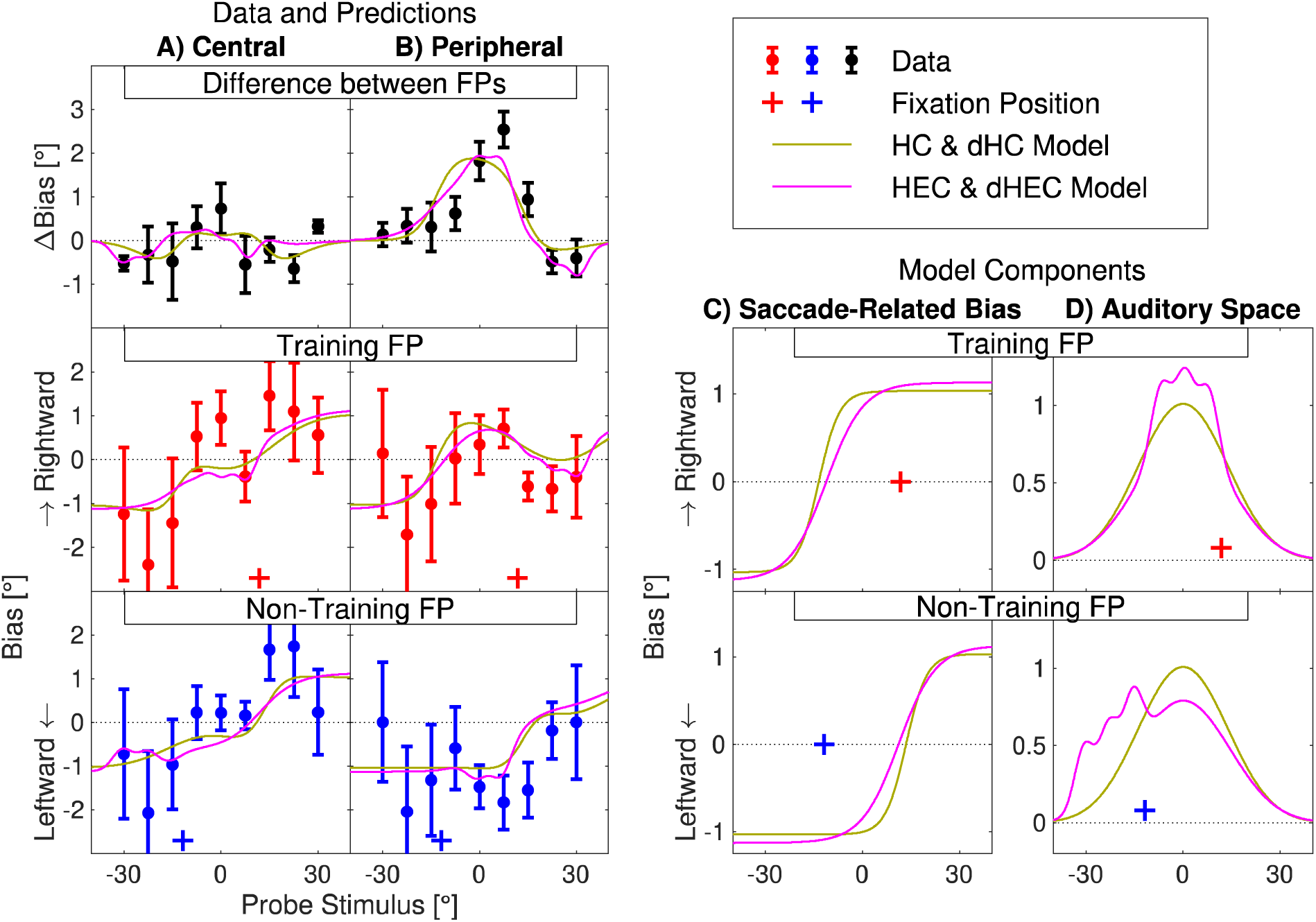
Model evaluation on No-Shift data. Model predictions (lines) and experimental data (symbols) for Central training experiment (A) and Peripheral training experiment (B). Middle and lower rows: Across-subject mean biases (± standard error) and model predictions for the two FPs separately. Upper row: Differences between the biases (± 1 standard error) for the two individual FPs and corresponding differences between the model predictions from the middle and lower rows. C) Predictions of the saccade-related bias model component of the HC and HEC models (dHC model performed identically to HC; dHEC identically to HEC). D) Predictions of the auditory space adaptation component for the central training experiment. Crosses: Fixation points.

Both models captured very well the difference data, which are near zero for the central training experiment and have a positive deviation for the peripheral training (upper row of panels A and B). This conclusion is confirmed by the AICc evaluation which showed no evidence that either of the models should be preferred even though the HEC model provides a slightly better fit (*ΔAIC* = 2.4 for HC). Also, the models captured all the important trends in the data for the two FPs (middle and bottom row). Specifically, for the central training data, they captured the slight expansion of the space for the central training data identical for both FPs (panel A), as well as the FP-dependence of the peripheral training data at the central locations (panel B).

A comparison of the error bars shows that there was a lot of across-subject variability when the individual FPs were considered (middle and bottom row), while a large portion of that variability was eliminated when the differences in biases across the FPs were computed (upper row). This illustrates why the models were fitted on the transformed data, as those were much more consistent across subjects, and, with the transformation, the fitting weighed the difference data (upper row) more, as they were much more reliable (FOOTNOTE 2).

Panels C and D show the predictions of the two model components (columns) separately for the two FPs (rows) for the central training data, such that the rows are aligned with the resulting central-data prediction in the same row of panel A (the peripheral predictions are based on the same model components except that the Auditory space prediction is shifted by 23.5° to the right from that in Figure 3D; not shown). Each prediction is, conceptually, a weighted sum of the two components (FOOTNOTE 3). The upper and lower panels of Figure 3C show that both models predicted similar saccade-related bias, consisting of expansion (bias toward periphery) at the peripheral target locations (+/-15°, +/- 22.5°, and +/-30°) and bias towards the fixation location for the central 3 locations (−7.5° to +7.5°). This saccade-related bias was then combined with the auditory space adaptation such that, at the training locations, the model predictions were shifted towards the AV response biases, which were near zero for both the central and peripheral training. The HC model predicts that this ventriloquism shift only occurred in HC RF (beige lines in the upper and lower panels of Figure 3D), while the HEC model predicts a considerable contribution of the EC RF (magenta lines at locations −30° to −15° in the bottom panel and at locations −7.5° to 7.5° in the top panel of Figure 3D). However, that contribution only had a small effect on the overall predictions because the difference between the AV responses and the saccade-related prediction, which scales the auditory space adaptation from panel D, was small for the no-shift data, resulting in the small differences between the beige and magenta lines in panel A.

### 5.2 Central data

Central Data simulation only fitted the central-training data from the AV-misaligned conditions (dashed lines in Figure 1C). For the mixed reference frame observed in these data, the simulation examined *whether the eye-referenced component is more consistent with the eye-referenced shift in adaptation region mechanism (HEC model) or the FP-dependent attenuation mechanism (dHC model)*. Figure 4A presents the results of this simulation. The top panel of Figure 4A shows the FP difference for the AV-misaligned data and the corresponding model predictions. The HC model’s prediction (beige) is fixed at zero, while the remaining three models fit the central-training data better, confirming that eye-referenced signals contribute to the ventriloquism adaptation in central region (the improvement vs. HC model in terms of AICc ranges from 5.9 to 15.6). Further, the HEC model’s AICc is worse by 9.6 compared to the dHC model, providing a strong evidence that the mixed reference frame observed behaviorally is driven by FP-dependent attenuation (dHC), not by ventriloquism signals in the EC RF (HEC). The HEC model (pink) underestimates the central data for targets at azimuths around 0° while it predicts a negative deviation at azimuths around −20°, not observed in the data, which the dHC (green) model does not predict. Finally, the dHC and dHEC models are comparable in terms of the AICs (difference of 1.4), while the dHEC model (purple) has the lowest MSE error, indicating that the EC-referenced shift in adaptation region mechanism might have a minor additional contribution to the adaptation effect.

**Figure 4:**
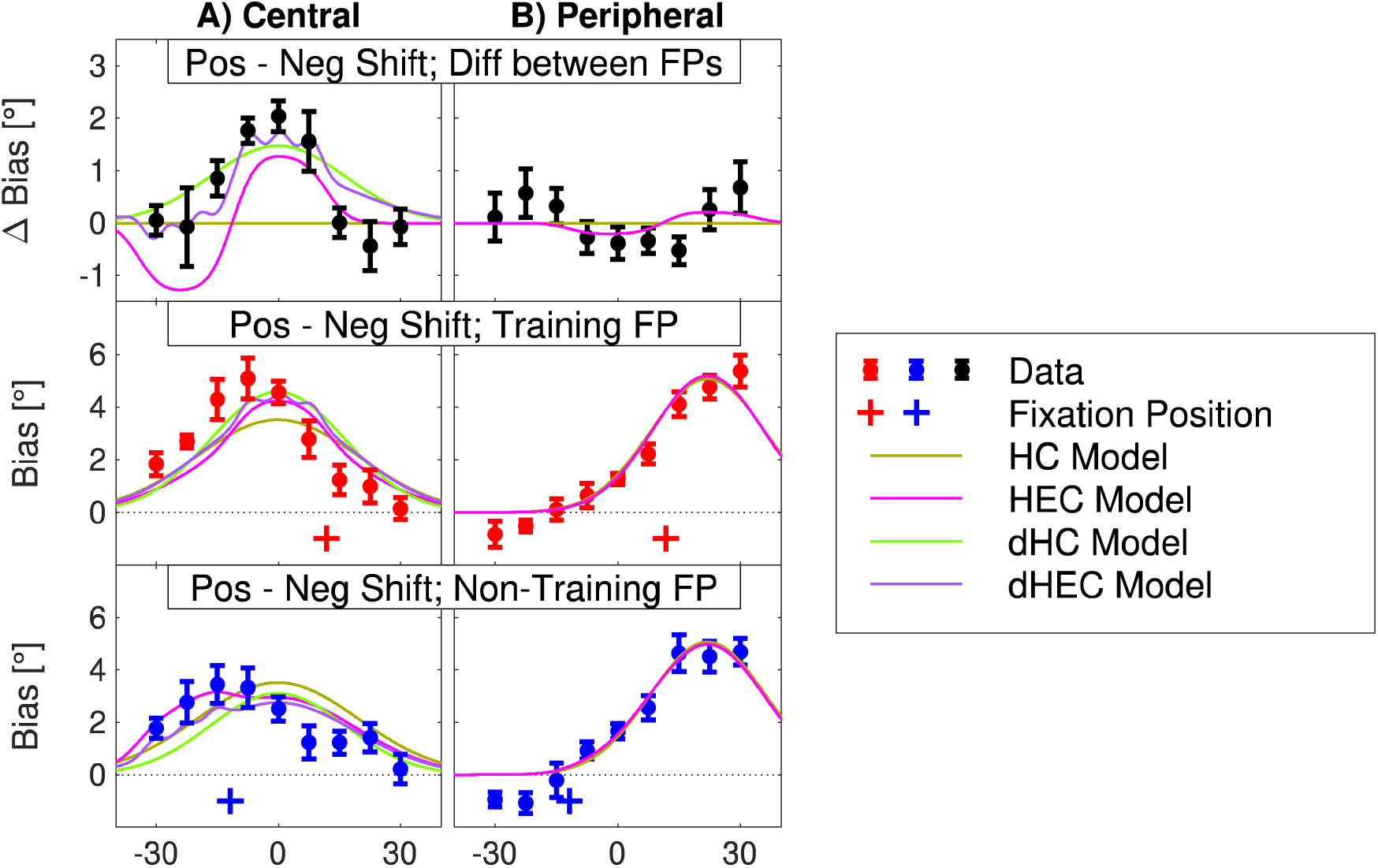
Model evaluations preformed separately on the AV-misaligned data from the Central training (A) and Peripheral training (B) experiments. The layout, color scheme and other aspects as in Figure 3A and 3B.

The middle and bottom panels of Fig 4A show the data and model predictions for the two FPs separately. All the models capture the basic profile of the adaptation. For the training FP (middle panel) the dHC model (green) is closer to the data than the HEC model (pink), in particular for the three left-most azimuths for which the difference prediction had the largest departure from the data (top panel). Interestingly, for the non-training FP (bottom panel), the HEC model captures the left-most triplet values better than the other two models. However, this improved non-training FP prediction results in the non-training FP values being larger than the corresponding training-FP values (pink line in the bottom panel is above the pink line in the middle panel), causing the difference prediction (top panel) to have the negative undershoot, not observed in the data.

The middle and lower panels also illustrate the functioning of the model for the AV-misaligned data for which the Auditory space representation component dominates the predictions. The dHC prediction (green line) in the lower panel is simply a scaled down version of the prediction from the middle panel, while the HEC prediction (pink) has two Gaussian components that are horizontally aligned and combined in the middle panel, while one of which is shifted to the left in the bottom panel. The dHEC model (purple) combines these two mechanisms to obtain predictions that tend to be closest to the data.

### 5.3 Peripheral data

Peripheral data simulation fitted only the peripheral-training data from the AV-misaligned conditions (solid lines in Figure 1C). The upper panel of Figure 4B shows that, in agreement with the data, all four models produced almost identical predictions of approximately 0-degree difference, confirming that the reference frame in this experiment was largely head-centered. Also, Table 2 shows that the HC model was indeed the best in terms of AICc and the EC-contribution parameters indicate a low contribution in the extended models (*d_f_* = 1 and *w_E_* = 0.04). The middle and bottom panels of Figure 4B show that the models captured the observed ventriloquism from individual FPs well, except for the two left-most points, for which the effect appears to be negative, which the current models cannot predict as Gaussians are used to describe the local character of the ventriloquism aftereffect.

### 6.4 Combined data

In the final evaluation, the four models were fitted to all data, combining the AV-aligned and AV-misaligned data from central and peripheral experiments (solid and dotted lines from Figure 1B and 1C). Figure 5 presents the results of this simulation in a layout similar to Figure 4, with the no-shift data added at the bottom. Overall, the model predictions in this simulation are less accurate than in the separate data set simulations (sections 6.1-6.3). The upper row of Figure 5 shows that the models tend to underestimate the FP-difference in the central data, in particular at azimuths near 0°, while they overestimate the FP-difference in the peripheral data, especially at the right-most azimuths. This pattern of results is caused by the largely linear operation of the model, which causes that the peripheral-data and central-data predictions are approximately identical when aligned with respect to the training region (i.e., the individual lines in the right-hand panel, when shifted to the left by 23.5°, are almost identical to the lines in the left-hand panel). Then, to minimize the error for both central and peripheral data, the model predictions are approximately in the middle of these two data sets. However, even with this constraint, the dHC and dHEC models perform significantly better than the HC and HEC model in terms of AICc (Table 2), again supporting the conclusion that the FP-dependent attenuation mechanism is the most likely mechanism causing the mixed reference frame observed for the central data, while the HEC mechanism only has a small contribution.

**Figure 5:**
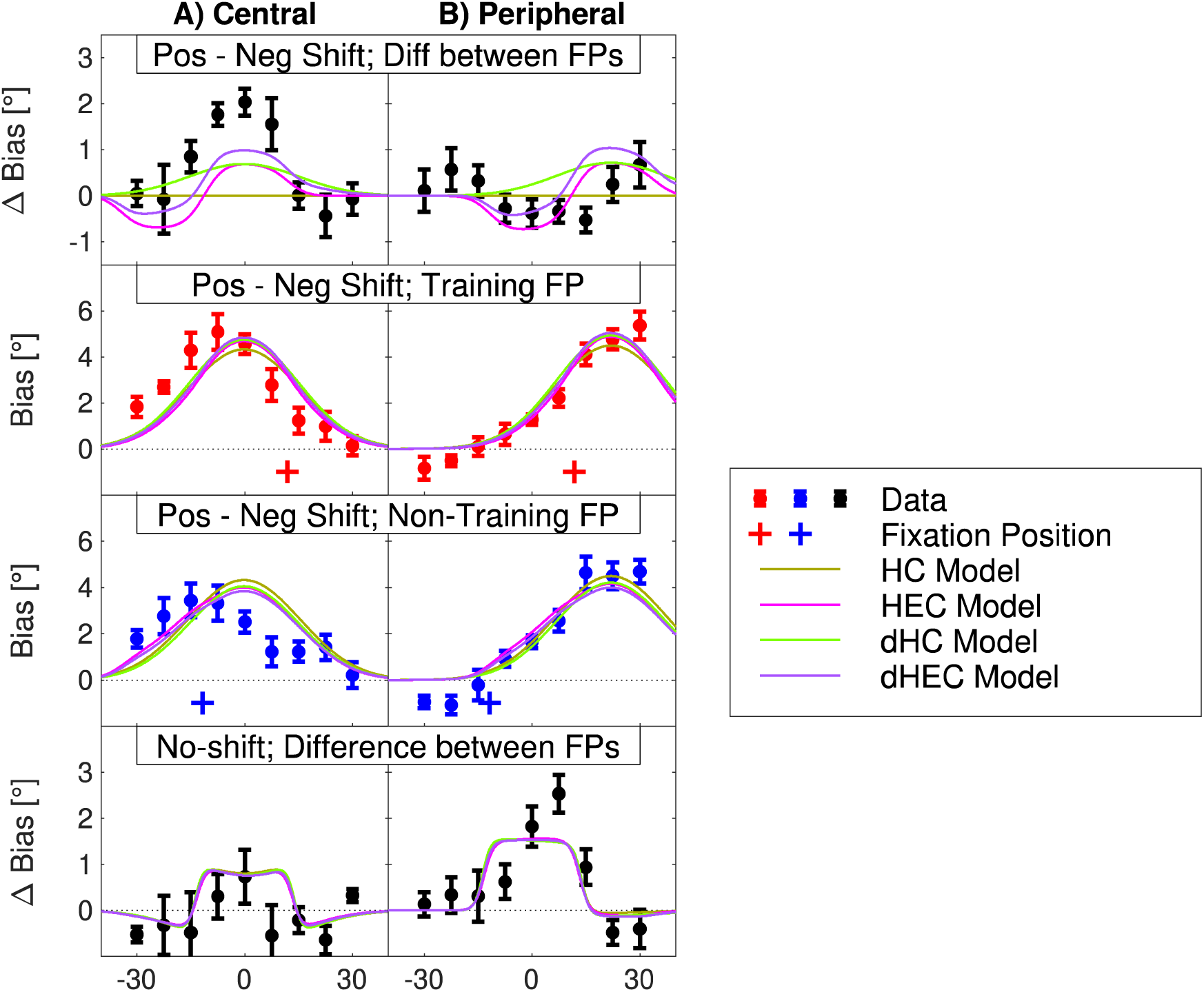
Model evaluation performed on the combined AV-aligned and AV-misaligned data from the Central training (A) and Peripheral training (B) experiments. The layout, color scheme and other aspects as in Figure 4A and 4B, with the no-shift difference data added to the bottom.

The two middle rows of Figure 5 show predictions separately for the two FPs. Overall, all the models predict the peripheral data well (right-hand column). On the other hand, the predictions for the central data show larger departures, especially for the middle targets and non-training FP, for which the bias in data is reduced more than the models predict. As discussed above, this is a limitation due to linear operation of the models.

Finally, the predictions for the AV-aligned data (bottom row) are again worse than when these data were considered separately (Figure 3), while the models do qualitatively capture that the biases are larger for the middle targets in the peripheral (panel B) than in the central (panel A) region. This less accurate prediction is again caused by linear combination of the saccade-related and auditory-space adaptation components in the model. Notably, all 4 models produce identical predictions of the AV-aligned data, since these predictions are mostly based on the saccade-related bias model component which is nearly identical across the 4 models.

## 6. Summary and Discussion

Our study introduced a model that combines ventriloquism-driven auditory space adaptation with saccade-related biases to predict the reference frame of the ventriloquism aftereffect. The model considers two forms of eye-centered signals influencing the auditory space representation: ventriloquism signals in eye-centered reference frame and FP-dependent attenuation. The main evaluation of the model, performed on the data from Kopco et al. (2009), found the FP-dependent attenuation mechanism to be the main eye-centered mechanism that caused the reference frame to be identified as mixed in that study. This result suggests that the auditory space representation is natively head-centered, and the FP-dependent signals only modulate the strength of the responses, thus making the result appear as mixed. Second, the model was also able to explain a new form of auditory space adaptation induced by AV-aligned stimuli in Kopco et al. (2019) by assuming a specific form of interaction between saccade-related bias and the ventriloquism adaptation. Finally, when the AV-aligned and AV-misaligned data from both studies were combined, the model predictions became less accurate, likely due to the limited linear interactions of the model components considered here. However, the model evaluation on the combined data still qualitatively supported the conclusions obtained in the separate evaluations.

The current model only uses the responses on AV training trials to predict the ventriloquism adaptation, independent of the size of the audio-visual disparity or of whether the disparity results in hypometric or hypermetric saccades. And it assumes that the ratio of observed ventriloquism aftereffect to the effect is constant, in our studies at approximately 0.5. With this simple assumption the model can also be applied to predict the results of other ventriloquism aftereffect studies, even those in which the ventriloquism effect was not measured, and those that did not use saccades for responding (in the latter case, the saccade-related bias component of the model can be simply omitted).

The model assumes that the ventriloquism aftereffect measured by saccades is influenced by some adaptive processes affecting the motor representations guiding the saccades to AV and/or auditory targets. This mechanism cannot be directly verified as currently available data are not consistent in terms of whether saccades to auditory targets are predominantly hypermetric or hypometric (Gabriel et al., 2010, Yao and Peck, 1997), while even less is known about saccades to misaligned AV targets. Additionally, eye-gaze-direction-dependent biases in sound localization have been previously observed even when saccades are not used for responding (Lewald and Ehrenstein, 1998, Razavi et al., 2007), and these are likely also influence the measured saccade biases. To tease these contributions apart, future studies need to assess the RF using response methods other than saccades (Kopco et al., 2015, Lewald and Ehrenstein, 1998).

The neural mechanisms of the ventriloquism aftereffect and its reference frame are not well understood. Cortical areas involved in ventriloquism aftereffect likely include Heschl’s gyrus, planum temporale, intraparietal sulcus, and inferior parietal lobule (Michalka et al., 2016, van der Heijden et al., 2019, Zatorre et al., 2002, Zierul et al., 2017). Multiple studies found some form of hybrid representation or mixed auditory and visual signals in several areas of the auditory pathway, including the inferior colliculus (Zwiers et al., 2004), primary auditory cortex (Werner-Reiss et al., 2003), the posterior parietal cortex (Duhamel et al., 1997, Mullette-Gillman et al., 2005, Mullette-Gillman et al., 2009), as well as in the areas responsible for planning saccades in the superior colliculus and the frontal eye fields (Schiller et al., 1979, Wallace and Stein, 1994). In the current model, the saccade-related component likely corresponds to the saccade-planning areas. The auditory space representation component likely corresponds to the primary or the higher auditory cortical areas, or the posterior parietal areas. There is growing evidence that, in mammals, auditory space is primarily encoded based on two or more spatial channels roughly aligned with the left and right hemifields of the horizontal plane (Dingle et al., 2012, Groh, 2014, Grothe et al., 2010, McAlpine et al., 2001, Salminen et al., 2009, Stecker et al., 2005). Considering such an extension might improve the model’s ability to predict the central and peripheral data simultaneously.

While most recalibration studies examined the aftereffect on the time scales of minutes (Radeau and Bertelson, 1974, Radeau and Bertelson, 1976, Recanzone, 1998, Woods and Recanzone, 2004), recent studies demonstrated that it can be elicited very rapidly, e.g., by a single trial with audio-visual conflict (Wozny and Shams, 2011). If it is the case that the adaptive processes underlying the ventriloquism aftereffect occur on multiple time scales, as also suggested in several models of slower ventriloquism aftereffect (Bosen et al., 2018, Watson et al., 2019), then an open question is whether the reference frame is the same at the different scales or whether it is different. The current results are mostly applicable to the slow adaptation on the time scale of minutes, while the RF on the shorter time scales has not been previously explored.

In summary, our study introduced a model of the reference frame of the ventriloquism aftereffect, while also considering the effect of saccade response biases on the measured reference frame. The model assumes simple linear combinations of model components to predict the data, which was sufficient to predict three data sets separately, but less so when predicting combined data. Future studies need to expand this model to be able to predict the combined data simultaneously, e.g., by consider non-linear combinations of the model components.

## Acknowledgments

This work was supported by the Slovak Scientific Grant Agency VEGA 1/0355/20 and by EU Danube Region Strategy grant ASH (Grant Nos. APVV DS-FR-19-0025, WTZ MULT 07/2020, 45268RE). The authors thank Piotr Majdak for his comments on an early version of this manuscript.

## Footnotes

**Footnote 1:** First, the data for the two training FPs were orthogonally transformed such that instead of using training and non-training FP, a sum and a difference across the two FPs was used. I.e., instead of having for each shift condition 18 data points corresponding to 9 locations at 2 FPs, we used 18 data points consisting of 9 locations summed across the two FPs and 9 locations for difference across the 2 FPs. Specifically, if *x_i,j,k_* represents a data point for the *i*-th target location (1 … 9), *j*-th AV-shift direction (P - positive, N - negative, 0 - no) and *k*-th fixation location (T – training, N – non-training), the first transformation produced a data set *y* with the same dimensionality as *x* where each data point *y_i,j,k_* is computed as:

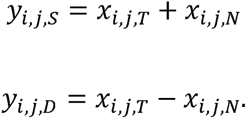

I.e., only the third dimension is changed such that the index *l* represents the operation performed across the fixations (S – sum; D – difference).

Second, the positive-shift and negative-shift condition data were transformed in a similar way, such that instead of positive and negative AV shift we used the aftereffect magnitude (i.e., a halved difference between the two shifts) and average across the two shifts. The no-shift data were left unmodified. Specifically, the second transformation produced a data set *z* with the same dimensionality as *y* where each data point *z_i,m,l_* is computed as:

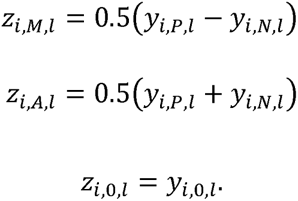

Here, only the second dimension is changed such that the index *m* represents the operation performed across the AV-shifts (M – magnitude of the aftereffect; A – average across positive & negative shifts; 0 – no shift). Note that the *z_i,M,l_* data are plotted as black lines in the bottom right panel of Figure 1C and the *z_i,_*_0*,l*_ data are plotted in the bottom panel of Figure 1B.

**Footnote 2:** Note that the second half of the transformed data set, the average across FPs, is not shown, as it can be easily visualized as an average of the individual FP data in the lower two rows of panel B.

**Footnote 3:** The prediction is obtained as a combination of the saccade-related bias and the auditory space component scaled 1) by the difference between the AV-training data and the saccade-related bias, and 2) by the weight parameter *w* (Eqs. 1 and 3). The auditory space graphs in Figure 3B are shown for the values of *d­* and [*r_AV_* – *r_E_*] fixed to 1 to illustrate the differences between the two models. However, actual values of the parameters need to be considered to generate the predictions in Figure 3A and 3B.

## References

Beierholm, U., Quartz, S. & Shams, L. 2009. Bayesian priors are encoded independently from likelihoods in human multisensory perception. Journal of Vision Arvo Journal, 9.

Bertelson, P., Frissen, I., Vroomen, J. & De Gelder, B. 2006. The aftereffects of ventriloquism: Patterns of spatial generalization. Perception and Psychophysics, 68, 428–436.

Bosen, A. K., Fleming, J. T., Allen, P. D., O’neil, W. E. & Paige, G. D. 2018. Multiple time scales of the ventriloquism aftereffect. PLoS ONE, 13.

Burnham, K. P. & Anderson, D. R. 2004. Multimodel Inference Understanding AIC and BIC in Model Selection. Sociological Methods & Research, 33, 261–304.

Carlile, S., Hyams, S. & Delaney, S. 2001. Systematic distortions of auditory space perception following prolonged exposure to broadband noise. Journal of the Acoustical Society of America, 110, 416–424.

Dingle, R. N., Hall, S. E. & Phillips, D. P. 2012. The three-channel model of sound localization mechanisms: interaural level differences. J Acoust Soc Am, 131, 4023–9.

Duhamel, J.-R., Bremmer, F., Benhamed, S. & Graf, W. 1997. Spatial invariance of visual receptive fields in parietal cortex neurons. Nature, 389, 845–848.

Gabriel, D. N., Munoz, D. P. & Boehnke, S. E. 2010. The eccentricity effect for auditory saccadic reaction times is independent of target frequency. Hearing Res, 262, 19–25.

Groh, J. M. 2014. Making space: how the brain knows where things are. Cambridge, MA: Harvard University Press.

Groh, J. M. & Sparks, D. L. 1992. Two models for transforming auditory signals from head-centered to eye-centered coordinates. Biological Cybernetics, 67, 291–302.

Grothe, B., Pecka, M. & Mcalpine, D. 2010. Mechanisms of sound localization in mammals. Physiol Rev 90, 983–1012.

Haessly, A., Sirosh, J. & Miikkulainen, R. A model of visually guided plasticity of the auditory spatial map in the barn owl. Seventeenth Annual Meetings of the Cognitive Science Society, 1995 Pittsburgh, PA. Erlbaum, 154–158.

Kopco, N., Lin, I. F., Shinn-Cunningham, B. G. & Groh, J. M. 2009. Reference Frame of the Ventriloquism Aftereffect. Journal of Neuroscience, 29, 13809–13814.

Kopco, N., Loksa, P., Lin, I. F., Groh, J. & Shinn-Cunningham, B. 2019. Hemisphere-specific properties of the ventriloquism aftereffect. J Acoust Soc Am, 146, EL177–183.

Kopco, N., Marcinek, L., Tomoriova, B. & Hladek, L. 2015. Contextual plasticity, top-down, and non-auditory factors in sound localization with a distractor. Journal of the Acoustical Society of America, 137.

Kording, K. P., Beacham, J. B., Ma, W. J., Quartz, S., Tenenbaum, J. B. & Shams, L. 2007. Causal Inference in Multisensory Perception. Plos One, 2.

Lewald, J. & Ehrenstein, W. H. 1998. Auditory-visual spatial integration: A new psychophysical approach using laser pointing to acoustic targets. Journal of the Acoustical Society of America, 104, 1586–1597.

Lingner, A., Pecka, M, Leibold, C., Grothe, B. 2018. A novel concept for dynamic adjustment of auditory space. Nature Scientific Reports, 8.

Mcalpine, D., Jiang, D. & Palmer, A. R. 2001. A neural code for low-frequency sound localization in mammals. Nature Neuroscience, 4, 396–401.

Michalka, S. W., Rosen, M. L., Kong, L., Shinn-Cunningham, B. & Somers, D. C. 2016. Auditory spatial coding flexibly recruits anterior, but not posterior, visuotopic parietal cortex. Cerebral Cortex, 26, 1302–1308.

Mullette-Gillman, O. A., Cohen, Y. E. & Groh, J. M. 2005. Eye-centered, head-centered, and complex coding of visual and auditory targets in the intraparietal sulcus. Journal of Neurophysiology, 94, 2331–2352.

Mullette-Gillman, O. A., Cohen, Y. E. & Groh, J. M. 2009. Motor-related signals in the intraparietal cortex encode locations in a hybrid, rather than eye-centered, reference frame. Cerebral Cortex, 19, 1761–75.

Odegaard, B., Ulrich, R., Carpenter, J. & Shams, L. 2019. Prior expectation of objects in space is dependent on the direction of gaze. Elsevier Cognition, 182, 220–226.

Odegaard, B., Wozny, D. R. & Shams, L. 2015. Biases in Visual, Auditory, and Audiovisual Perception of Space. PLOS Computational Biology, 11, e1004649.

Oess, T., Ernst, M. O. & Neumann, H. 2020. Computational principles of neural adaptation for binaural signal integration. PLOS Comput Biol, 16.

Pouget, A., Deneve, S. & Duhamel, J. R. 2002. A computational perspective on the neural basis of multisensory spatial representations. Nature Reviews Neuroscience, 3, 741–747.

Radeau, M. & Bertelson, P. 1974. The after-effects of ventriloquism. Quarterly Journal of Experimental Psychology, 26, 63–71.

Radeau, M. & Bertelson, P. 1976. The effect of a textured visual field on modality dominance in a ventriloquism situation. Perception and Psychophysics, 20, 227–235.

Razavi, B., O’neill, W. E. & Paige, G. D. 2007. Auditory Spatial Perception Dynamically Realigns with Changing Eye Position. Journal of Neuroscience, 27, 10249–10258.

Recanzone, G. H. 1998. Rapidly induced auditory plasticity: The ventriloquism aftereffect. Proceedings of the National Academy of Sciences of the United States of America, 95, 869–875.

Salminen, N. H., May, P. J., Alku, P. & Tiitinen, H. 2009. A population rate code of auditory space in the human cortex. PLoS One 4:e7600.

Shinn-Cunningham, B. G., Kopco, N. & Martin, T. J. 2005. Localizing nearby sound sources in a classroom: Binaural room impulse resonses. Journal of the Acoustical Society of America, 117, 3100–3115.

Schiller, P. H., True, S. D. & Conway, J. L. 1979. The effects of frontal eye field and superior colliculus ablations on eye movement. Science, 206, 590–592.

Stecker, G. C., Harrington, I. A. & Middlebrooks, J. C. 2005. Location Coding by Opponent Neural Populations in the Auditory Cortex. PLoS Biology, 3, e78.

Taboga, M. 2017. Normal distribution - Maximum Likelihood Estimation. Lectures on probability theory and mathematical statistics, Third edition. Kindle Direct Publishing.

Van Der Heijden, K., Rauschecker, J. P., De Gelder, B. & Formisano, E. 2019. Cortical mechanisms of spatial hearing. Nature Reviews Neuroscience, 20, 609–623.

Van Opstal, J. 2016. The Auditory System and Human Sound-Localization Behavior, Elsevier.

Wallace, M. T. & Stein, B. E. 1994. Cross-modal synthesis in midbrain depends on input from cortex. Journal of Neurophysiology, 71, 429–432.

Watson, D. M., Akeroyd, M. A., Roach, N. W. & S., W. B. 2019. Distinct mechanisms govern recalibration to audio-visual discrepancies in remote and recent history. Sci Rep, 9.

Watson, D. M., Akeroyd, M. A., Roach,, N. W., Webb B. S. 2021. Multiple spatial reference frames underpin perceptual recalibration to audio-visual discrepancies. Plos One, 16.

Werner-Reiss, U., Kelly, K. A., Trause, A. S. & Underhill, A. M. 2003. Eye position affects activity in primary auditory cortex of primates. Current Biology, 13, 554–562.

Woods, T. M. & Recanzone, G. H. 2004. Visually Induced Plasticity of Auditory Spatial Perception in Macaques. Current Biology, 14, 1559–1564.

Wozny, D. R., Beierholm, U. & Shams, L. 2010. Probability Matching as a Computational Strategy Used in Perception. PLoS Comput Biol, 6.

Wozny, D. R. & Shams, L. 2011. Recalibration of Auditory Space following Milliseconds of Cross-Modal Discrepancy. Journal of Neuroscience, 31, 4607–4612.

Yao, L. & Peck, C. K. 1997. Saccadic eye movements to visual and auditory targets. Exp Brain Res, 115, 25–34.

Zatorre, R. J., Bouffard, M., Ahad, P. & Belin, P. 2002. Where is ‘where’ in the human auditory cortex? Nat Neurosci, 5, 905–9.

Zierul, B., Roder, B., Tempelmann, C., Bruns, P. & Noesselt, T. 2017. The role of auditory cortex in the spatial ventriloquism aftereffect. Neuroimage, 162, 257–268.

Zwiers, M. P., Van Opstal, A. J. & Paige, G. D. 2003. Plasticity in human sound localization induced by compressed spatial vision. Nat Neurosci, 6, 175–81.

Zwiers, M. P., Versnel, H. & Van Opstal, A. J. 2004. Involvement of monkey inferior colliculus in spatial hearing. J Neurosci, 24, 4145–56.

